# STAT pathway activation limits the Ascl1-mediated chromatin remodeling required for neural regeneration from Müller glia in adult mouse retina

**DOI:** 10.1101/753483

**Authors:** Nikolas L. Jorstad, Matthew S. Wilken, Levi Todd, Paul Nakamura, Nick Radulovich, Marcus J. Hooper, Alex Chitsazan, Brent A. Wilkerson, Fred Rieke, Thomas A. Reh

## Abstract

Müller glia can serve as a source for retinal regeneration in some non-mammalian vertebrates. Recently we found that this process can be induced in mouse Müller glia after injury, by combining transgenic expression of the proneural transcription factor Ascl1 and the HDAC inhibitor TSA. However, new neurons are only generated from a subset of Müller glia in this model, and identifying factors that limit Ascl1-mediated MG reprogramming could potentially make this process more efficient, and potentially useful clinically. One factor that limits neurogenesis in some non-mammalian vertebrates is the STAT pathway activation that occurs in Müller glia in response to injury. In this report, we tested whether injury induced STAT activation hampers the ability of Ascl1 to reprogram Müller glia into retinal neurons. Using a STAT inhibitor, in combination with our previously described reprogramming paradigm, we found a large increase in the ability of Müller glia to generate neurons, similar to those we described previously. Single-cell RNA-seq showed that the progenitor-like cells derived from Ascl1-expressing Müller glia have a higher level of STAT signaling than those that become neurons. Using Ascl1 ChIP-seq and DNase-seq, we found that developmentally inappropriate Ascl1 binding sites (that were unique to the overexpression context) had enrichment for the STAT binding motif. This study provides evidence that STAT pathway activation reduces the efficiency of Ascl1-mediated reprogramming in Müller glia, potentially by directing Ascl1 to inappropriate targets.

## INTRODUCTION

Functional regeneration of retinal neurons occurs naturally in teleost fish and many amphibians^1–6^. In zebrafish, the Müller glia (MG) respond to a variety of injury models by generating cells that resemble multipotent progenitor cells found in the developing retina. These MG-derived progenitors have the capacity to produce all types of retinal neurons and restore visual function^7^. In amphibians and embryonic birds, the pigmented epithelial cells undergo a similar transition to retinal progenitor cells, and can regenerate a new, laminated retina^8, 9^.

In adult birds and mammals, functional regeneration does not occur spontaneously after retinal injury. Neurotoxic damage to retinal neurons in newly hatched chicks causes the MG to undergo the initial stages of the process that occurs in fish, but few of the MG-derived progenitors go on to make neurons and it is not known whether the few regenerated MG-derived neurons can functionally integrate into the existing retinal circuitry^1^. Injury to the mammalian retina has been studied most extensively in rodents, and as in the bird, retinal injury does not initiate a spontaneous regenerative response^10^. Attempts to stimulate MG proliferation after injury by stimulating specific signaling pathways with growth factors and small molecules have led to some evidence for new neurogenesis^11^; however, none of these treatments have been sufficient to regenerate functional neurons from MG in mice^12–14^.

Recently, we have found that transgenic overexpression of the proneural bHLH Ascl1 enables MG to generate functional neurons in mice. We found that in mice up to two weeks old, Ascl1 alone can induce neurogenesis from MG after N-Methyl-D-aspartic acid (NMDA) excitotoxic damage^15^. More recently, we demonstrated that Ascl1 and the HDAC inhibitor trichostatin A (TSA) were together sufficient to induce MG to regenerate functional neurons after retinal injury in adult mice^16^. While these studies were encouraging and demonstrated for the first time that new neurons generated in adult mice can be integrated into the mature retinal circuit, the majority of the MG in treated retinas did not undergo neurogenesis.

In the course of our analysis of single-cell RNA-seq data from the previous study, we noticed that those MG that failed to reprogram to neurons had a high level of STAT3 signaling. MG from fish, birds and mammals all respond to retinal damage by rapidly activating STAT3 signaling^17–20^. In the mouse retina, STAT3 signaling in MG is associated with reactive “gliosis”; however, in the fish retina, STAT3 is required for damage-induced MG-mediated neuronal regeneration, and JAK/STAT activation is sufficient to induce retinal regeneration in the absence of injury^17, 21, 22^. In the chick retina, however, experiments have suggested that STAT signaling instead may limit regeneration: inhibition of STAT3 increases the neurogenesis from MG in damaged retinas^20^. Since an increase in STAT signaling is well known to occur in glia after neuronal injury, we tested whether the injury response was potentially reducing Ascl1-mediated reprogramming. We tested this hypothesis by inhibition of STAT signaling, and found that a STAT inhibitor, in combination with our previously described treatment paradigm, doubled the efficiency of neuron regeneration in vivo. Analysis of ChIP-seq data of Ascl1-expressing MG suggests that STAT signaling may act to redirect Ascl1 to inappropriate regulatory sites in the genome, and implicate the Id1 and Id3 inhibitors of bHLH factors in this process. Our results extend previous work in fish and birds on the importance of JAK/STAT signaling in retinal regeneration. Together with the results from the non-mammalian vertebrates, it appears that an initial activation of the STAT pathway may be necessary to start the regeneration process, but sustained activation of the pathway may then become limiting.

## RESULTS

### STAT pathway inhibition improves MG-derived neurogenesis in vivo

To test whether the activation of the STAT pathway in MG reduces their ability to be effectively reprogrammed by Ascl1, we inhibited this pathway in an experimental system in which Ascl1 can be targeted specifically to MG. We previously described^15, 16^ a MG-specific tamoxifen inducible mouse model of Ascl1-overexpression (*Glast*-CreER:Rosa-flox-stop-*LNL*-tTA:*tetO*-mAscl1-ires-GFP). Adult mice received up to five consecutive daily intraperitoneal (IP) injections of tamoxifen (1) to induce Ascl1 expression in the MG, followed by (2) an intravitreal injection of NMDA to induce degeneration of inner retinal neurons, and then (3) an intravitreal injection of either TSA or TSA and the potent STAT inhibitor, SH-4-54 (referred to as ANT and ANTSi treatment, henceforth) (Figure 1A). SH-4-54 successfully blocks the phosphorylation of STAT3’s tyrosine 705 residue in Ascl1-overexpressing MG and was not found to be toxic at the concentrations used for this study (Figure S1). As previously described, ANT treatment resulted in GFP+ MG-derived neurons that expressed bipolar neuron genes Otx2 and Cabp5, in addition to many MG that failed to undergo neurogenesis (Figure 1B). When the STAT pathway inhibitor was co-injected with the TSA (ANTSi treatment), there was a striking increase in the number of MG-derived neurons (Figure 1C, 1D). Quantification of the mature bipolar gene Cabp5 revealed a more than two-fold increase in the number of MG-derived neurons (24 ± 5% and 52 ± 3%, n = 6 and n = 8, for ANT and ANTSi treatments, respectively; Figure 1E-F). All Cabp5+ GFP+ cells also expressed Otx2 and all Cabp5+ cells had neuronal morphology; however, not all Otx2+ cells expressed Cabp5 or had neuronal morphology, as previously described^16^.

**Figure 1.**
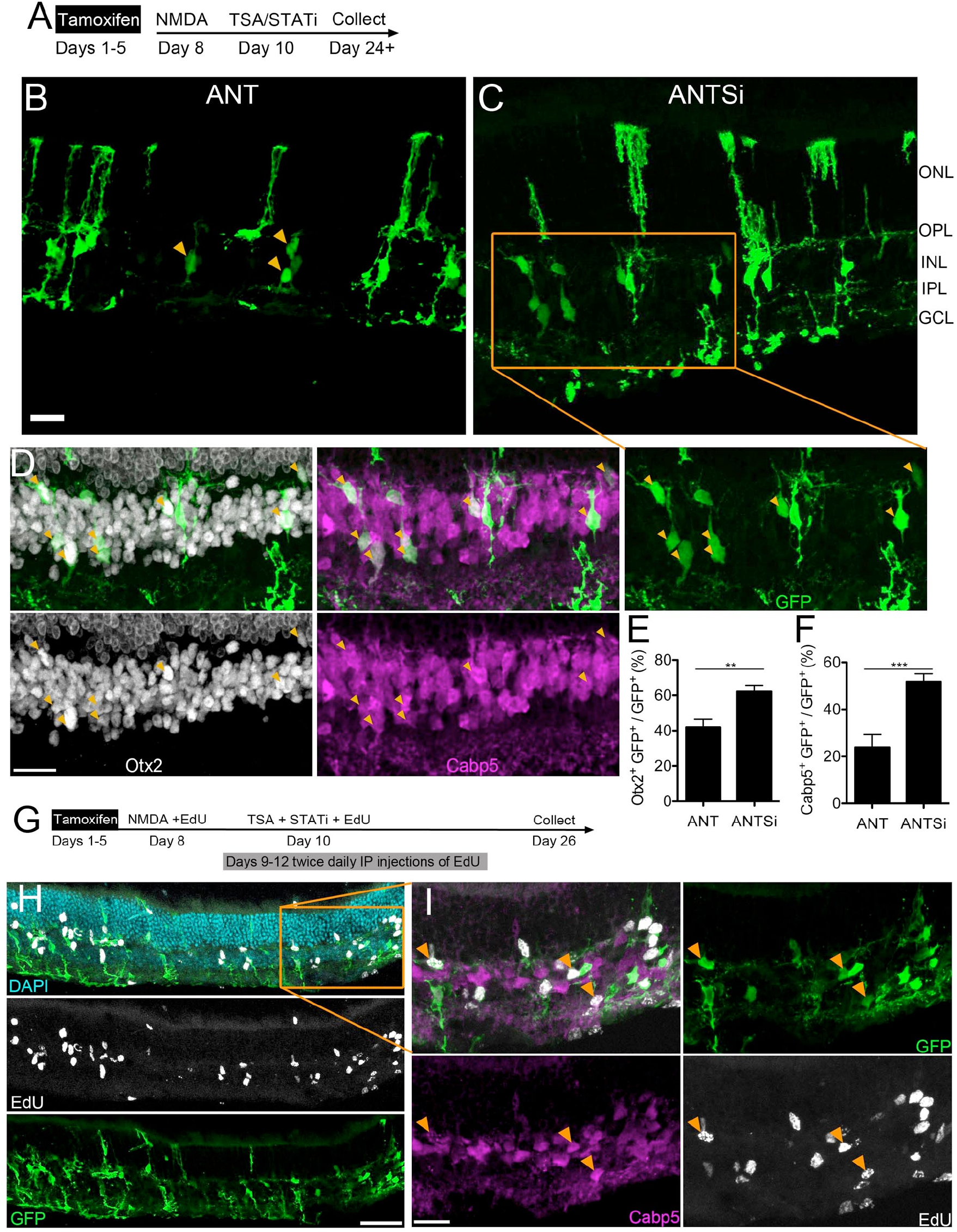
STAT pathway inhibition increases number of Müller glial-derived neurons. **A)** Experimental paradigm for increasing MG-derived regeneration efficiency. Tamoxifen is administered for up to 5 consecutive days, followed by NMDA damage a few days after tamoxifen, followed by administration of TSA and/or STAT inhibition a couple days after damage. Retinas were collected a minimum of two weeks after TSA/STATi. **B)** Representative image showing ANT-treated adult retina with MG-derived neurons. **C)** Representative image showing ANTSi-treated adult retina with increased number of MG-derived neurons. **D)** Shows enlargement of ANTSi-treated retinas from **C**. Orange arrows indicate Cabp5+ Otx2+ GFP+ cells. All images are flattened Z-stacks. Scale bars for **B-C** are 20 µm. ONL = Outer Nuclear Layer, OPL = Outer Plexiform Layer, INL = Inner Nuclear Layer, IPL = Inner Plexiform Layer, GCL = Ganglion Cell Layer. **E)** Quantification of Otx2 in ANT (n = 16) and ANTSi-treated (n = 13) retinas. **F)** Quantification of Cabp5 in ANT (n = 6) and ANTSi-treated (n = 8) retinas. ANT vs ANTSi treatments in **E** and **F** were significantly different by unpaired *t*-test at **P=0.0023 and ***P=0.0006, respectively. **G)** Experimental paradigm for testing where ANTSi-treated MG proliferate prior to neurogenesis. **H)** Representative image from proliferation experiment showing EdU and GFP colocalization. **I)** Shows enlargement of **H** and highlights GFP+ EdU+ Cabp5+ MG-derived neurons (orange arrows). Scale bars for **H-I** are 50 µm and 20 µm, respectively.

To determine whether the MG-derived neurons arise from mitotic proliferation or directly differentiate into neurons we intravitreally co-injected EdU with the NMDA on treatment day 8 and also with the TSA and STATi on treatment day 10 (Figure 1G). Additionally, we performed twice daily IP injections of EdU from treatment day 9 through 12. After collecting the retinas on treatment day 26, we found that many of the MG and MG-derived neurons were labeled with EdU (Figure 1H). Additionally, most of the MG-derived neurons labeled with EdU, and Cabp5, and had neuronal morphology (Figure 1I, orange arrows). These results show that many of the new neurons regenerated in the ANTSi condition are the result of mitotic proliferation of the MG. We found that approximately 53% of GFP+ cells in the ANTSi condition were co-labeled with EdU.

### ANTSi treated MG-derived neurons make synaptic connections with existing circuitry

In our previous study, we found that MG-derived neurons generated by ANT treatment express synaptic proteins, make connections with cone terminals, and successfully integrate into the existing retinal circuitry and functionally respond to light stimuli^16^. In this study, we found that similar to ANT-treated neurons, ANTSi-treated MG-derived neurons express synapse proteins Ctbp2 and Psd95. Labeling retinal sections from the ANTSi treatment group for cone blue opsin (Opn1sw) showed MG-derived neurons contacting cone photoreceptors terminals in the OPL, indicative of photoreceptor-bipolar network connections (Figure 2A-C).

**Figure 2.**
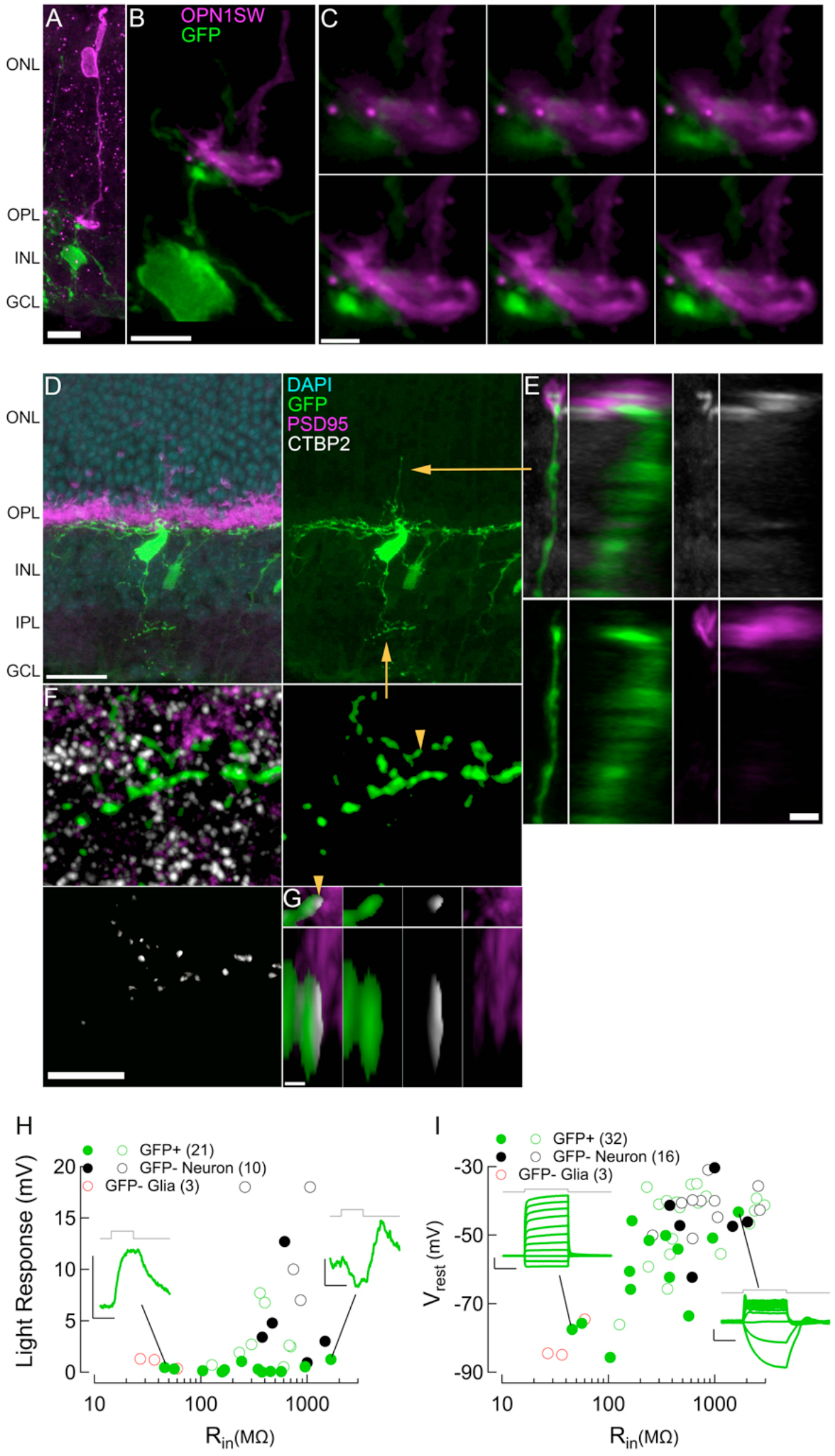
ANTSi-treated MG-derived neurons integrate into existing retinal circuits. **A and B)** Example images of MG-derived neuron (green) contacting cone photoreceptors (magenta) in the OPL, scale bars are 10 µm and 5 µm, respectively. **C)** Enlargement of **B** showing 0.18 µm z-stack steps of the MG-derived neurons making contact with cone pedicle, scale bar is 2 µm. **D)** Example image of MG-derived neuron with neuronal process in OPL and IPL, scale bar is 20 µm. **E)** Enlargement of yellow arrow from **D** showing a Psd95+ Ctbp2+ photoreceptor synapse onto an apical MG-derived neuronal process in ONL. Left image sets show XY projection and right image sets show YZ projection, scale bar is 2 µm. **F)** Enlargement of yellow arrow from **D** showing Ctbp2 staining within the MG-derived neuronal processes in IPL. Upper right image shows stringent GFP mask used to identify Ctbp2 puncta within MG-derived neurons, scale bar is 5 µm. **G)** Enlargement of yellow arrowhead from **F** showing Ctbp2 within masked GFP process, directly apposed to Psd95 staining, consistent with synaptic specializations. Upper image sets show XY projection and lower image sets show XZ projection, scale bar is 0.5 µm. **H)** Population data for input resistance and visual responses recorded with the current-clamp technique. **I)** Population data for input resistance and resting membrane potential measurements. **H** and **I**, Solid colored circles indicate new measurements recorded from ANTSi treatment condition and hollow colored circles indicate measurements from ANT treatment condition from a previous study^16^. Green traces in **H** show maximum evoked light response from 500-ms luminance stimulation for MG (left) and MG-derived neuron (right). Green traces in **I** show representative responses to equal increasing steps of injected current for MG (left) and MG-derived neuron (right). ANTSi-treated recordings were performed at 2, 5, and 7 weeks post-TSA and STATi administration.

To determine if ANTSi-treated MG-derived neurons express synaptic proteins in the correct cellular locations and form synaptic specializations with other cell types, we stained for Ctbp2 and Psd95 (Figure 2D-G). In the OPL, we found Psd95+ Ctbp2+ photoreceptor synapses onto MG-derived neuron processes, indicative of a photoreceptor-to-MG-derived neuron synaptic specialization. Interestingly, we also found these contacts in the ONL (Figure 2D-E) with photoreceptors synapsing onto the apical process of MG-derived neurons. This irregular location for photoreceptor synapse formation is likely due to the retraction of photoreceptor terminals^23^. In the IPL, we found Ctbp2 puncta within the MG-derived neuron terminals (Figure 2F-G). Pre-synaptic Ctbp2 within the MG-derived neurons was directly apposed to Psd95 from post-synaptic cell partners, indicative of a MG-derived neuron output onto other neurons in the IPL (Figure 2G). Our data suggest that MG-derived neurons make synaptic specializations with photoreceptors in the OPL and post-synaptic cells in the IPL, consistent with our previous findings that MG-derived neurons are able to make synaptic connections within the existing retinal circuitry.

Next, we performed whole-cell electrophysiology recordings from GFP+ and GFP-cells to determine if ANTSi-treated MG-derived neurons have electrical properties consistent with retinal neurons. We plotted the population data from recorded cells onto a plot of previously recorded ANT-treated cells from another study^16^ (Figure 2H-I). We found that GFP+ cells had much higher input resistance (*R*_in_) than GFP-MG and the resting membrane potentials (*V*_rest_) of GFP+ cells were depolarized compared to the GFP-MG. Most GFP+ cells exhibited *R*_in_ and *V*_rest_ properties similar to GFP-neurons (Figure 2I), suggesting that the GFP+ cells’ electrical properties were indeed more similar to retinal neurons. To determine if GFP+ cells integrated into the retinal circuitry and received synaptic input from photoreceptors responding to light stimulus, we exposed dark adapted ANTSi-treated retinas to incremental increases of luminance and recorded from MG-derived neurons (Figure 2H). Similar to our previous study of ANT-treated MG-derived neurons^16^, we found examples of ANTSi-treated MG-derived neurons responding to light increments by rapidly depolarizing, as would be expected for ON-preferring retinal neurons. Light responses presented with amplitudes ∼2 mV, which were not as robust as some of the cell recordings from our previous study; however, these recordings were performed on cells after 2, 5, and 7 weeks post-treatment, whereas our previous study included cells from later time points ranging from 4-15 weeks. Taken together, these findings indicate that MG-derived neurons in the ANTSi treatment condition have retinal neuron electrical properties and receive synaptic input from photoreceptors resulting in light-evoked visual responses.

### Single-cell RNA-seq on reprogrammed Müller glia

To further characterize the effects of STAT inhibition during MG reprogramming, we ran single-cell RNA-seq (scRNA-seq) on FACS-purified MG and MG-derived cells 14 days after ANTSi treatment, similar to that described in an earlier study^16^. In total 648, 823, and 2283 cells were analyzed from WT, ANT, and ANTSi treatments, respectively. All three (WT, ANT, and ANTSi) datasets were combined, normalized, scaled and analyzed in R with Seurat and Monocle^24–26^. Projecting all three treatments onto a single tSNE plot shows ten distinct clusters (Figure 3A), differentially associated with the different treatment conditions (Figure 3C). We identified the cell types comprising the clusters by their unique expression of specific known marker genes. Some of these genes are shown projected onto the individual clusters (Figure 3B) and additional genes are shown associated with the clusters as a heatmap (Figure 3F). For example, Clusters 3 and 4 were comprised of cells almost exclusively from the untreated (WT) condition and expressed genes normally present in MG, such as Glul and Aqp4; these were classified as “MG” clusters (Figure 3D). Clusters 0 and 2 showed enrichment for progenitor genes (Dll1; Figure 3B) but lacked neuronal gene expression (Otx2 and Cabp5) and showed reduced glial gene expression (Glul). We therefore classified clusters 0 and 2 as “Progenitor-like” cells. Clusters 1 and 6 showed enrichment for neuronal genes and greatly reduced expression of glial genes and were classified as “MG-derived Neurons”. In addition to these major clusters, there were two contaminating cell populations that we do not think were derived from MG. These include microglial cells, identified by their expression of Aif1 (IBA1), and a small number of cells that express Rho and other photoreceptor genes that we consider contaminating “Rods”. These two populations are present in the FACS-purified cells regardless of treatment group, whereas the progenitor-like cells and MG-derived neurons are only present in the ANT and ANTSi conditions (Figure 3C-D). Clusters 8 and 9 contained a small number of cells and were not enriched for any particular retinal cell type (Figure 3D; “Other”).

**Figure 3.**
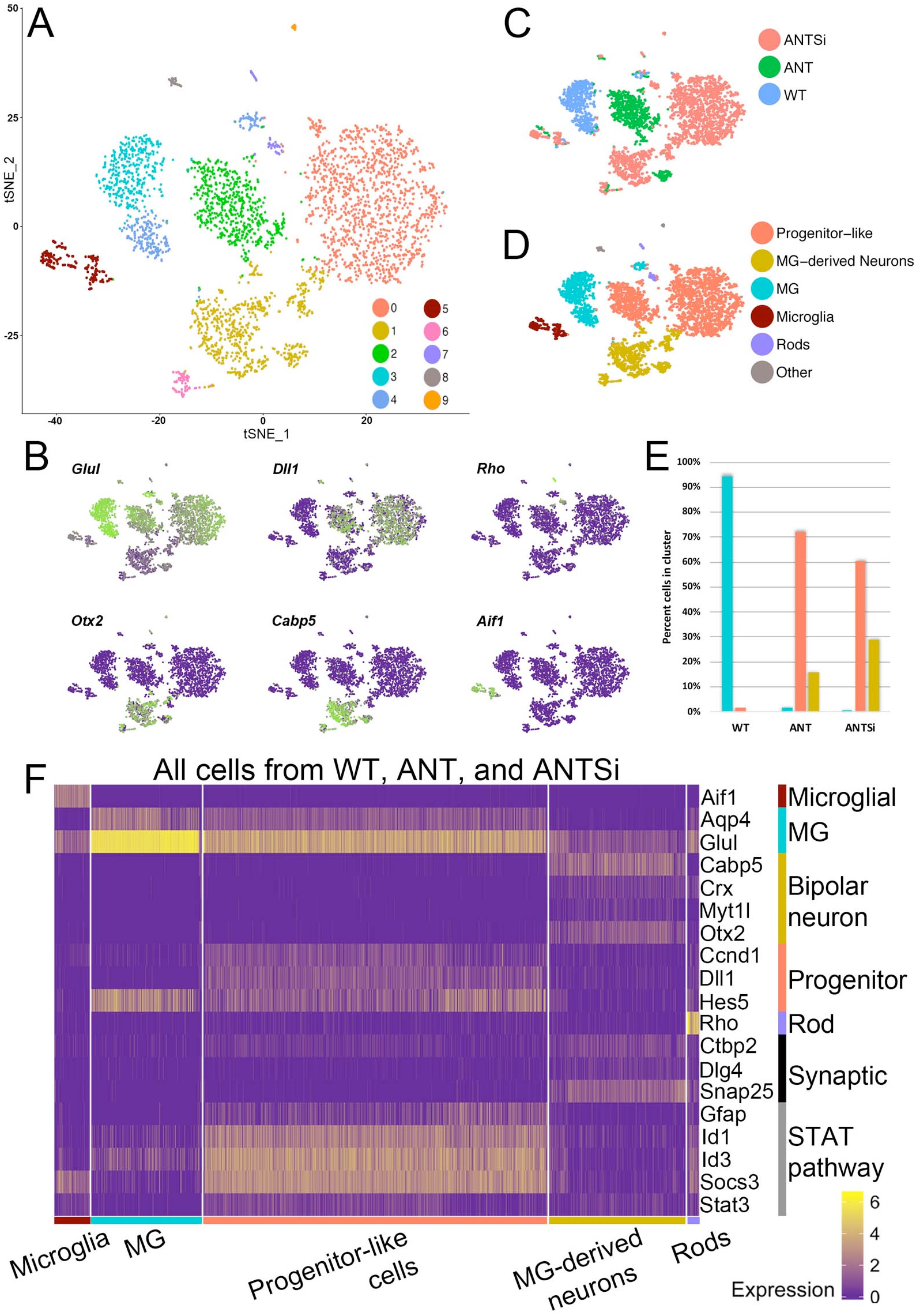
ANTSi treatment results in more MG-derived neurons by scRNA-seq. **A)** A tSNE plot of FACS-purified WT MG and ANT-treated cells from out previous study^16^, and ANTSi-treated cells. **B)** Feature plots of glial (Glul), progenitor (Dll1), neuronal (Cabp5 and Otx2), rod (Rho), and microglial (Aif1) gene expression to identify clusters by cell type. Green shows high-expressing, purple shows non-expressing cells. **C)** Plot from **A** colored by treatment condition. **D)** Clusters pseudo colored by cell type as determined by **C**. Microglia and rods made up contaminating populations of ∼5% and 2% of the total cells in each treatment group, respectively. **E)** Graph showing the fraction of each treatment that was comprised of MG, Progenitor-like cells, and MG-derived neurons. ANTSi treatment increases the population of MG-derived neurons and decreases the number of progenitor-like cells relative to ANT treatment. **F)** Heatmap showing all cells from all three treatments. MG-derived neurons express synaptic genes and do not express Stat pathway targets. Progenitor-like cells highly express Stat pathway targets. Scale shows log2 expression.

The immunofluorescent analysis of retinas from the ANTSi condition indicated that a greater percentage of progenitor-like cells differentiate into neurons when compared with the ANT treatment. We analyzed the scRNA-seq results to determine whether a similar trend could be detected. Quantifying the number of WT, ANT, and ANTSi cells in the MG, progenitor-like, or MG-derived neuron clusters (Figure 3E) revealed nearly double the number of MG-derived neurons in the ANTSi treatment compared to ANT (29% of total ANTSi cells vs. 16% of total ANT cells). The increase in the MG-derived neuron population in the ANTSi treatment was accompanied by a decrease in the number of progenitor-like cells. These data support the immunofluorescent analysis that STAT inhibition results in a population shift of MG from progenitor-like cells to MG-derived neurons and doubles reprogramming efficacy.

To better understand the trajectory of the reprogramming process, we used the Monocle analysis package on a subset of the data that only included MG, progenitor-like cells, and MG-derived neurons from clusters 0, 1, 2, 3, 4, and 6 (Figure 4A). We found the pseudotime scale tracked with our classified cell types, with MG being the root state and transitioning through the progenitor-like cell state and eventually progressing towards MG-derived neurons (Figure 4B). We observed cells decreasing expression of glial genes (e.g. Glul) as they progress towards progenitor-like cells and MG-derived neurons (Figures 4C-D). Cells in the progenitor-like clusters exclusively express progenitor genes, such as Dll1 and as cells move towards the MG-derived neuronal clusters they upregulate neuronal genes, like Cabp5.

**Figure 4.**
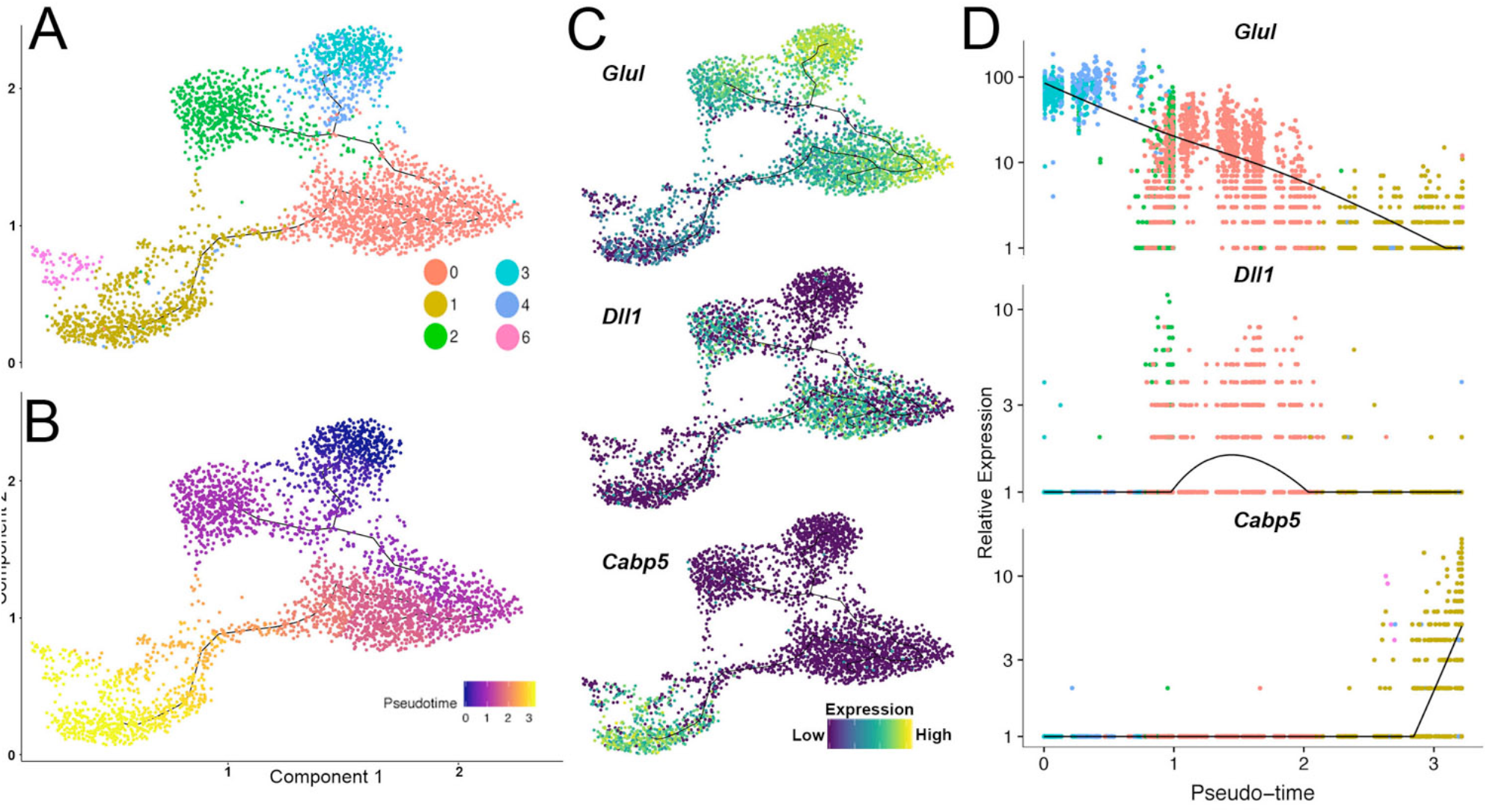
Pseudotime analysis of scRNA-seq datasets. **A)** Trajectory analysis in Monocle showing clusters from subset of data that includes MG, Progenitor-like cells, and MG-derived neurons. **B)** Trajectory analysis showing pseudotime progression, with cluster 3 (from **A**) being the root state. **C, D)** Gene expression of Glul, Dll1, and Cabp5 shown as trajectory plot **C**, and pseudotime plot **D**.

Although the trajectory analysis shows that MG progress to neurons through a progenitor-like state, it also suggests that this state is not identical in the ANT and ANTSi conditions. The progenitor-like cells in these two conditions cluster separately and although both clusters express progenitor genes, like Dll1, the pseudotime plot suggests ANT reprogrammed cells are “closer” to MG than the cells from the ANTSi treated retinas. This led us to hypothesize that progenitor cells may be kept in a stable, but non-neural state, by STAT pathway activation. Indeed, the progenitor-like cells highly express various STAT pathway target genes (e.g. Gfap, Id1, Id3, Socs3, Figure 3F), while MG and MG-derived neurons have reduced or no expression of STAT targets.

### ChIP-seq for Ascl1 in P0 mouse retinal progenitors and reprogrammed Müller glia

To explore potential mechanisms by which STAT inhibition promotes Ascl1-mediated retinal regeneration, we carried out ChIP-seq for Ascl1 in MG and compared this with retinal progenitors. In order to determine the endogenous binding pattern of Ascl1 in retinal progenitor cells on a genome-wide scale, we performed ChIP-seq for Ascl1 in P0 retina (Figure 5A) and identified 22,251 peaks using HOMER^27^ with a False Discovery Rate (FDR) of 0.1%. The top-scoring motif in these peaks (using MEME) was the canonical Ascl1 E-Box (Figure S2A)^28^. The great majority of the Ascl1 peaks overlapped with an accessible chromatin region (Figure 5; P0 DNase Track)^29^. We compared the retinal Ascl1 peaks to previous ChIP-seq for Ascl1 in other neural tissues. For this analysis, we focused on the highest scoring peaks that replicated in two independent P0 retinal Ascl1-ChIP-seq runs (7587 sites, Figure 5A). When the three-different neural Ascl1-ChIP-seq datasets were compared, we found that the majority of the Ascl1-bound regions in each type of neural tissue were unique to that tissue (Figure 5B); for the peaks present in retina and another neural tissue, there was approximately the same degree of overlap for each of the combinations, while only a relatively small subset of peaks were common to all neural tissues (764; Figure 5A-B; Core Ascl1 track). Gene Ontology analysis using the GREAT algorithm^30^ showed that those peaks unique to retina were enriched for eye, optic and retinal GO terms (for mouse phenotype), while those regions common to all three neural tissues instead have enrichment for GO terms of non-retinal neural tissues (e.g. telencephalon; Figure 5B). In addition, the genes associated with these GO terms for the retinal-specific regions were genes expressed more highly in retina than in other regions of the CNS, like cerebral cortex or spinal cord (e.g. Rax, Six3, Atoh7, Otx2; Figure 5B), while those genes associated with the Ascl1-bound regions common to all three neural tissues were potentially representative of a more generic neural program of differentiation (Notch1, Insm1, and Dll1), though it is important to note that there are also region-specific regulatory sites for common targets, like Hes5 and Sox2.

**Figure 5.**
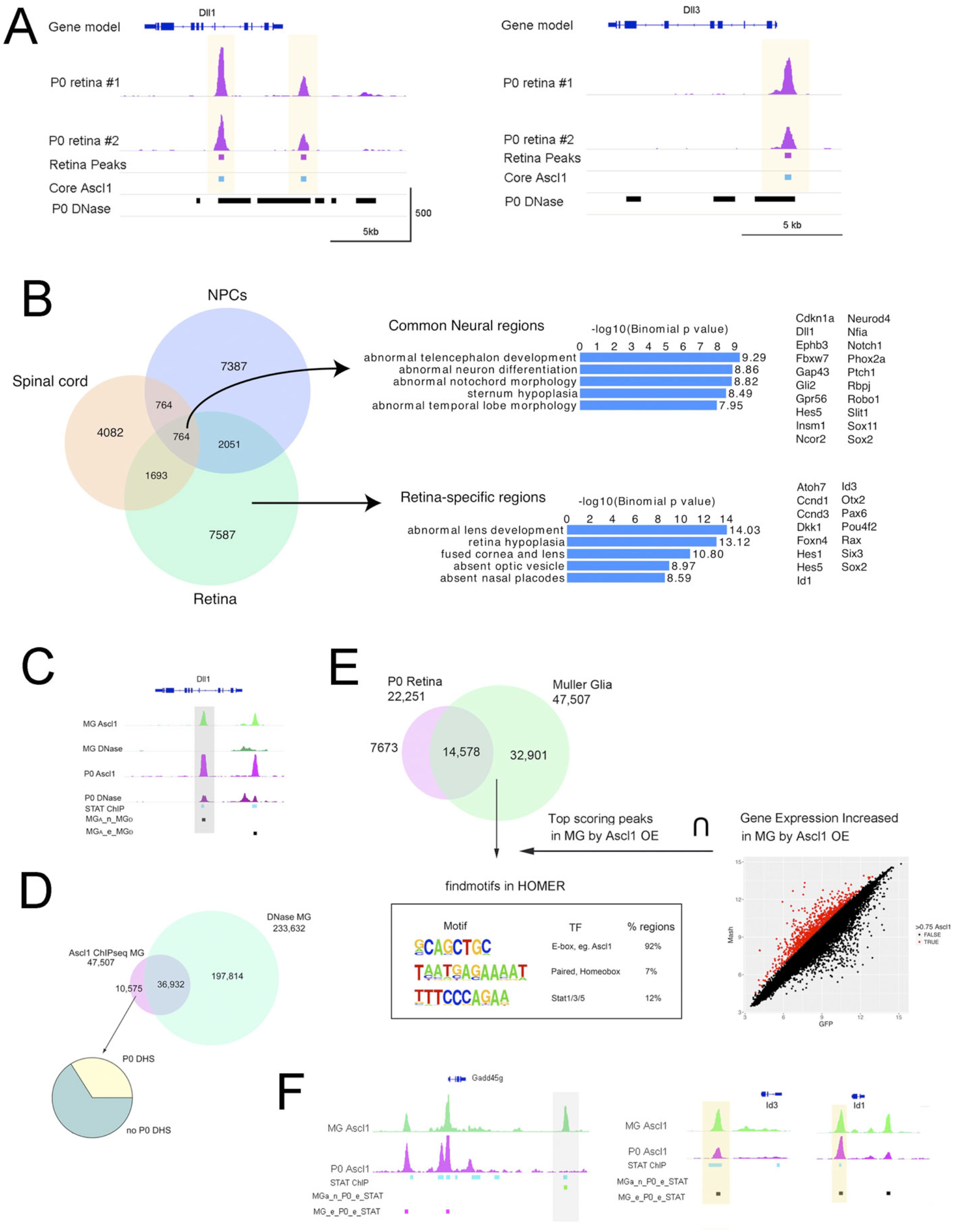
Ascl1 ChIP-seq from Ascl1-overexpressing MG and P0 retinal progenitors. **A)** Epigenetic analyses of progenitor genes Dll1 and Dll3. Tracks show biological replicate Ascl1 ChIP-seq peaks from P0 whole retina (P0 retina #1, #2), bars indicate peaks that were called from peak-calling algorithm HOMER with FDR of 0.1% (Retina Peaks track), peaks that were called from retinal, spinal cord, and NPC Ascl1 ChIP-seq datasets (Core Ascl1 track), and DNase-seq peaks from P0 whole retina showing accessible chromatin (P0 DNase track). Yellow highlights indicate core/common binding sites. Scale at the bottom track (X-axis = kilobases of genomic DNA, Y-axis = reads per million, RPM). **B)** Venn diagram showing proportions of overlap for Ascl1 ChIP-seq peaks between P0 retina, NPCs, and spinal cord (Left). Gene Ontology analysis of retinal-specific and core/common Ascl1 ChIP-seq peaks using GREAT algorithm and example genes in these categories (Right). **C)** Epigenetic comparison of Ascl1-overexpressing MG (MG Ascl1 and MG DNase tracks) with P0 developing retina (P0 Ascl1 and P0 DNase tracks). Additional tracks showing previously described Stat3 ChIP-seq peaks from brain oligodendrocytes (STAT ChIP) and comparative peak overlap analyses of Ascl1 peaks without DHSs (MG_A_n__MG_D_) or with DHSs (MG_A_e__MG_D_). **D)** Above, Venn diagram showing proportions of overlap of Ascl1 ChIP-seq peaks with DNase-seq peaks from Ascl1-overexpressing MG. Below, Pie chart showing the proportion of Ascl1 ChIP-seq peaks that have a P0 DNase peak present. **E)** Venn diagram showing proportions of overlap from Ascl1 ChIP-seq peaks between P0 developing retina and Ascl1-overexpressing MG (Top). Integrative analysis looking for motifs that were enriched at Ascl1-overexpressing MG-specific Ascl1 peaks (developmentally inappropriate), are located within ±5kb of the T.S.S., and are associated with 0.75 increase in gene expression (from a previous Ascl1-virus vs GFP-virus microarray). Top scoring motifs meeting these criteria are presented in box (E-box 92%, Paired homebox 7%, Stat1/3/5 12%) (Below). **F)** Epigenetic comparison of Gadd45g, Id1, and Id3 gene loci in Ascl1-overexpressing MG (MG Ascl1) and P0 developing retina (P0 Ascl1). Additional tracks showing previously described Stat3 ChIP-seq peaks (STAT ChIP), and comparative peak overlap analyses of sites containing a MG Ascl1 peak, a P0 Ascl1 peak, and a Stat3 peak (MG_A_e__P0__e__STAT) or sites containing a MG Ascl1 peak, a Stat3 peak, but no P0 Ascl1 peak (MG_A_n__P0__e__STAT). Yellow highlights indicate strong Ascl1 binding sites during development and forced Ascl1 expression, Grey highlights indicate anomalies.

To determine the extent to which Ascl1 binds to similar regions in MG as in retinal progenitors, we performed ChIP-seq for Ascl1 in MG during the reprogramming process. Retinas from P12 mice (germline-rtTA: *tetO*-Ascl1) were dissociated and the MG were grown in culture. After 7 days, Ascl1 expression was induced in the purified MG by addition of doxycycline to the culture medium. Chromatin was then collected after 6 days post-Ascl1 induction and subsequently processed for Ascl1 ChIP-seq. The subsequent sequencing reads were filtered and mapped to the mouse genome (Figure 5C) and peaks called using HOMER. In total, 47,507 peaks were called with a False Discovery Rate (FDR) of 0.1%; MEME analysis shows that the top-scoring motif was the canonical Ascl1 E-Box, similar to that described above for the P0 retina (Figure S2A).

We performed a binding site overlap analysis between the peaks from the two cell populations. As shown in Figure 5E, 31% (14,578/47,507) of Ascl1 binding regions in MG overlap an ‘appropriate’ Ascl1 binding site (i.e. one present in P0 retinal progenitor cells); however, the majority (∼70%) of the Ascl1 bound regions in MG are not present in the P0 retina. This discrepancy is not simply due to the higher number of binding sites in the MG, since 34% (7,673/22,251) of retinal progenitor Ascl1 binding sites are not bound in the MG. Therefore, although the majority of P0 retinal progenitor Ascl1 bound regions are also bound by Ascl1 in MG, there is substantial binding of Ascl1 to “inappropriate” sites in the genome of MG. In addition, there are also many potential cis-regulatory sites in progenitors that are not bound in the MG.

When we further analyzed the Ascl1 binding sites that are specific to retinal progenitors and compared them with those bound in the MG using the ‘GREAT’ gene ontology algorithm, the majority of top enriched terms related to neurogenesis (e.g. – ‘neural retina development’, ‘layer formation in cerebral cortex’), suggesting that these Ascl1 binding sites are important for normal neuronal development of the retina and not dispensable binding events (Figure S2B). Ascl1 binding sites that occurred in both progenitors and MG were enriched for neurogenic and gliogenic terms (e.g. – ‘negative regulation of gliogenesis’, ‘negative regulation of oligodendrocyte differentiation’) (Figure S2B). However, the majority of Ascl1 binding sites specific to MG were not associated with retinal development (e.g. – ‘filopodium assembly’, ‘regulation of mitochondrial membrane permeability’) (Figure S2B). Therefore, while Ascl1 binds 66% of all appropriate sites in retinal progenitors, the majority of binding sites in MG are inappropriate and potentially not productive towards neurogenic reprogramming.

Studies in induced pluripotent cell reprogramming have shown that induction of pluripotency genes by Yamanaka factors^31^ (Oct4, Sox2, cMyc, and Klf4) is initially limited by the accessible chromatin sites in the somatic cell genome already bound by existing transcription factors^32^. We wondered whether the inappropriate sites bound by Ascl1 in MG are due to opportunistic binding to DNase-Hypersensitive sites (DHSs) that are present in the MG and not in the retinal progenitors. Therefore, we quantified the overlap between Ascl1 ChIP-seq peaks and DHSs that are present in MG. As shown in Figure 5D, 78% of Ascl1 binding sites occurred within DNase-hotspots in the MG and 22% occurred within non-hypersensitive chromatin. This result suggests that while most of the binding in MG occurs at sites that are already accessible, between one fifth and one quarter of the Ascl1-bound sites in MG were “pioneered” by Ascl1, consistent with prior results in other cells^33^. We hypothesized that these pioneer Ascl1 binding events in MG were actually occurring at regions of DNase-hypersensitivity in the P0 retina. This scenario would be productive towards the goal of reprogramming the cis-regulatory landscape of MG towards that of retinal progenitors. Therefore, we quantified the overlap between pioneer Ascl1 binding sites in MG and P0 DNase-hotspots (Figure 5D). We found that 34% (3591/10,575) of the pioneered sites overlap a P0 DNase-Hotspot. However, a much greater number, 66% (6984/10,575) of the pioneered sites, do not overlap a DNase-Hotspot in either P0 retina or MG, and thus are potentially non-productive pioneered sites.

The preceding analysis suggests that while a third of Ascl1 pioneering events are productive for reprogramming towards a retinal progenitor state, a majority of pioneering events appear to be unproductive towards reprogramming to the retinal progenitor state. We hypothesized that other transcription factors present in the MG might be collaborating with Ascl1 to direct it to these inappropriate sites^32^. To determine which factors might be associated with the inappropriate Ascl1 ChIP-seq peaks in MG (i.e. not present in progenitors), we used HOMER to identify sequence motifs enriched in these regions (for genes that increase >0.75 fold in reprogrammed MG, over P0 progenitors from our previous microarray^34^). We found that Ascl1 was the top site followed by a generic homeodomain motif, but these regions were also significantly enriched for the STAT consensus site (Figure 5E). Thus, the activation of the STAT pathway in the MG after retinal injury may direct Ascl1 to inappropriate sites on the DNA and thereby reduce its ability to reprogram MG more fully to progenitors/neurons.

Among the most highly upregulated genes in the Ascl1-reprogrammed progenitor-like cells include the Inhibitor of Differentiation genes (Id1, Id2, and Id3). These genes code for proteins that are similar to bHLH transcription factors, like Ascl1, except they lack a DNA-binding domain, and thus can form heterodimers with the bHLH family of transcription factors and prevent their ability to activate transcription^35, 36^. The scRNA-seq analysis shows that the Ascl1-overexpressing MG that failed to convert to neurons (and retained RPC-like gene expression) highly expressed Id genes Id1, Id2, and Id3 (Figure 3F). Ascl1 ChIP-seq and DNase-seq from both P0 and MG show strong binding sites at accessible chromatin at both Id1 and Id3 promoters (Figure 5F). Additionally, a previously generated STAT3 ChIP-seq dataset from oligodendrocytes in the brain (GEO: GSM2650746) indicated STAT3 binding sites in the same region as the Ascl1 binding sites (STAT ChIP track, Figure 5F). Taken together, these findings suggest that STAT pathway activation and/or Ascl1 binding may induce Id gene expression, which inhibits Ascl1 from initiating neurogenesis in a subset of MG.

### ChIP-seq for Ascl1 in control and STATi-treated reprogrammed Müller glia

To directly assess the effect that STAT pathway activation has on Ascl1 binding at Id genes and more broadly across the genome during MG reprogramming, we performed an Ascl1 ChIP-seq on control and STATi-treated MG. Frozen stocks of MG from P12 germline-rtTA: *tetO*-Ascl1 mice were thawed and cultured to confluency for 2 days in high FBS growth media (Figure 6A). Cells were then given a low FBS media with STATi or vehicle for the remaining 4 DIV. At 3 DIV (1 day after STATi), doxycyline was administered to induce Ascl1 overexpression in the MG. Cells were then collected and fixed for ChIP-seq after 6 DIV. FASTQ files were aligned to the mm10 genome using Bowtie2 and peaks were called using MACS2. Representative tracks show Ascl1 Chip-seq for the Ascl1 expressing MG and sister cells treated with STATi are shown in Figure 6B. We observed 106,063 and 89,793 high confidence peaks in the Ascl1 control and Ascl1/STATi-treated samples, respectively (Figure S3A). The majority of peaks overlapped between the two samples, with 80,169 sites being common to both treatment conditions. Both datasets contained the canonical Ascl1 E-box as the top scoring motif, as determined by HOMER, with strong central enrichment of peaks from both datasets around the Ascl1 motif (Figure S3B).

**Figure 6.**
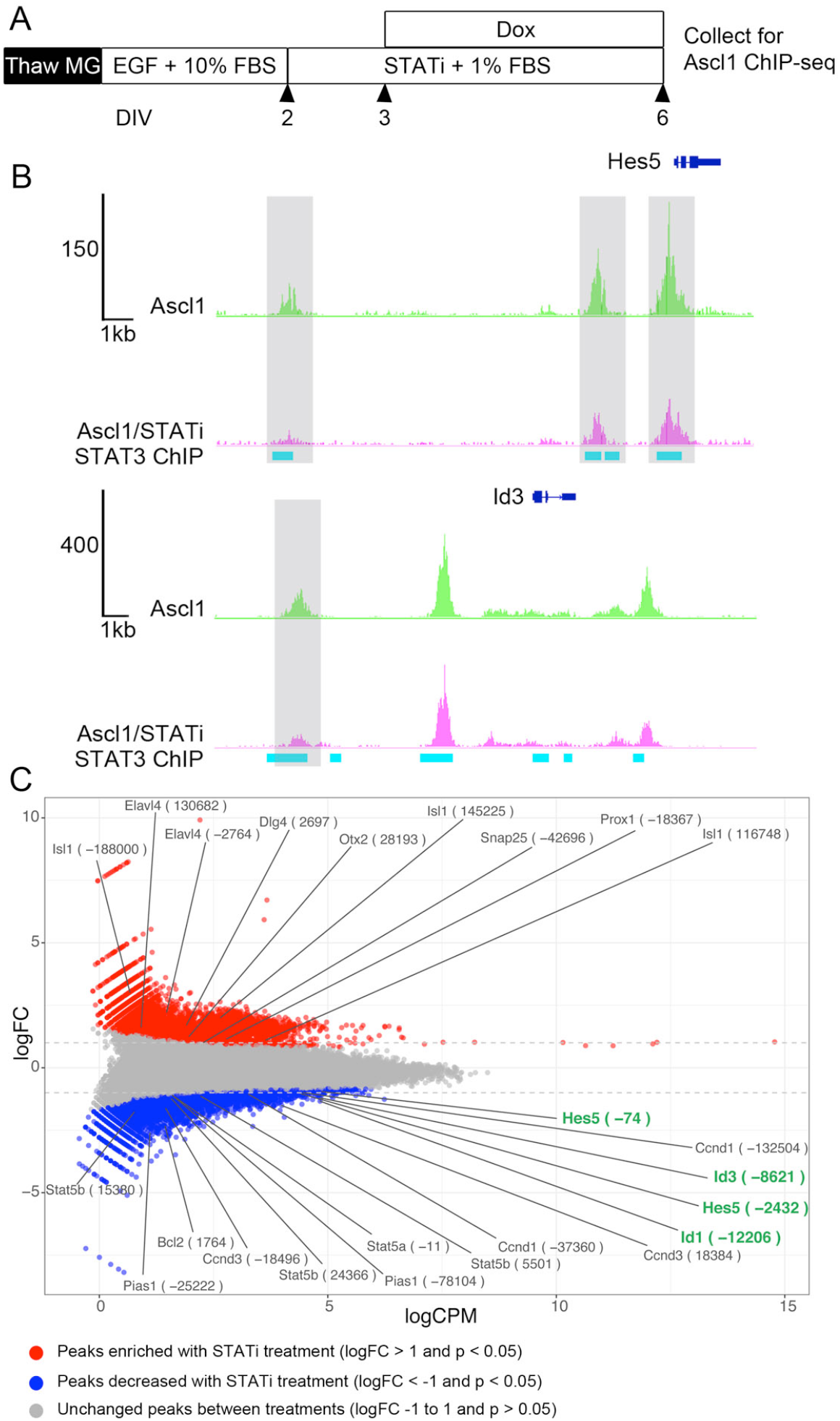
Ascl1 ChIP-seq from Ascl1-overexpressing MG with or without STATi. **A)** Experimental paradigm for Ascl1 ChIP-seq. **B)** Representative tracks of Ascl1 ChIP-seq datasets from Ascl1 and Ascl1/STATi conditions, and STAT3 ChIP-seq data from oligodendrocytes (GEO: GSM2650746). Gray highlights show peaks that were significantly decreased in STATi-treated cells. **C)** MA plot of all Ascl1 and Ascl1/STATi-treated Ascl1 ChIP-seq peaks. Red shows peaks that were significantly enriched in the STATi-treated cells with a logFC >1 and blue shows peaks that were significantly decreased in the STATi-treated cells with a log FC <-1.

Based on our findings that inappropriate Ascl1 binding sites were frequently found with the STAT motif, we hypothesized that inhibition of the STAT signal would reduce Ascl1 recruitment to these sites. To identify genes that might be regulated by STAT signaling in the MG during reprogramming with Ascl1, we performed a differential analysis between the control Ascl1 and Ascl1/STATi datasets using edgeR^37, 38^. Individual peaks were compared between the two datasets and plotted as a MA plot in Figure 6C. Ascl1 ChIP-seq peaks that were significantly reduced in the STATi treatment condition (Figure 6C, blue points) were present at various STAT pathway target genes (e.g. Bcl2, Pias1, Stat5a/b), and Id1 and Id3 were among these. Figure 6B shows decreased Ascl1 binding at STAT3 bound regions near Id1 and Id3 after STATi treatment, with gray highlights indicating the corresponding (significantly reduced) peaks from Figure 6C. Together, these findings add further support to the hypothesis that STAT signaling may redirect Ascl1 to non-productive cis-regulatory regions, and in some cases, inhibitors of neural reprogramming, like Id1, Id3 and Hes5.

In addition to reducing the binding of Ascl1 to STAT target regions in MG, inhibition of the STAT pathway leads to increases in Ascl1 binding to non-STAT targets. Ascl1 ChIP-seq peaks that were significantly enriched in the STATi treatment condition (Figure 6C, red points) were present at neuronal genes associated with amacrine and bipolar cell fates (e.g. Elavl4, Otx2, Isl1, and Prox1) as well as synaptic genes (e.g. Dlg4 and Snap25), consistent with our observations that this treatment leads to an increase in neuronal differentiation. By contrast, peaks located near genes associated with progenitor cells were also significantly decreased in the STATi treatment condition (e.g. Ccnd1 and Ccnd3). Thus, STATi results in enrichment of Ascl1 binding at neuronal genes, suggesting that inhibiting STAT pathway activation during reprogramming leads to more productive binding of Ascl1 at genes necessary for neurogenesis.

### Id genes are dysregulated in the presence of Ascl1

To determine the effects of Ascl1 overexpression and damage on STAT pathway activation and Id1 expression, we performed a NMDA time course (Figure 7A). Under normal conditions, in WT animals Id1 is not present at detectable levels in MG (Figure S4A). At 1 day and 2 days post-NMDA, Id1 is highly expressed in MG but begins to decline at 4 days and returns to basal levels by 9 days post-NMDA (Figure S4B-F). In the Ascl1-overexpressing mice, all MG express Id1 at similarly high levels by 2 days post-NMDA; both the Ascl1-overexpressing GFP+ MG, and the non-Ascl1-overexpressing GFP-Sox2+ MG (Figure 7B). By 4 days post-NMDA damage, Id1 was found to be at lower levels in the GFP-MG and highly expressed in the Ascl1 GFP+ MG (Figure 7C). Eleven days after TSA treatment, the GFP+ cells that show signs of neuronal morphology and downregulation of Sox2 have basal levels of Id1 expression (Figure 7F, orange arrows). By 14 days post-NMDA damage, high Id1 expression became restricted to the MG that express Ascl1 (the GFP+ MG) (Figure 7D). Additionally, we performed RT-qPCR for Id1 on whole retinas taken from undamaged WT and Ascl1 overexpressing retinas as well as NMDA treated retinas (Figure 7E, n = 4 mice per group). Ascl1 overexpressing retinas treated with NMDA expressed significantly more Id1 at 4 days post-NMDA than all other treatment groups. Interestingly, the Ascl1-overexpressing retinas have similar levels of Id1 at baseline as the WT mice treated with NMDA. Taken together, these findings suggest that the normally transiently activated target genes of STAT become constitutively active in the presence of Ascl1 overexpression. Constitutive expression of ID proteins in Ascl1 expressing MG likely results in the ID proteins binding Ascl1, antagonizing the pro-neurogenic effects of Ascl1^35, 36^. Further evidence supporting this hypothesis is the fact that ANT-treated MG-derived neurons do not have any detectable Id1 expression (Figure 7F, orange arrows), while the adjacent non-reprogrammed MG do have Id1 expression (white arrow).

**Figure 7.**
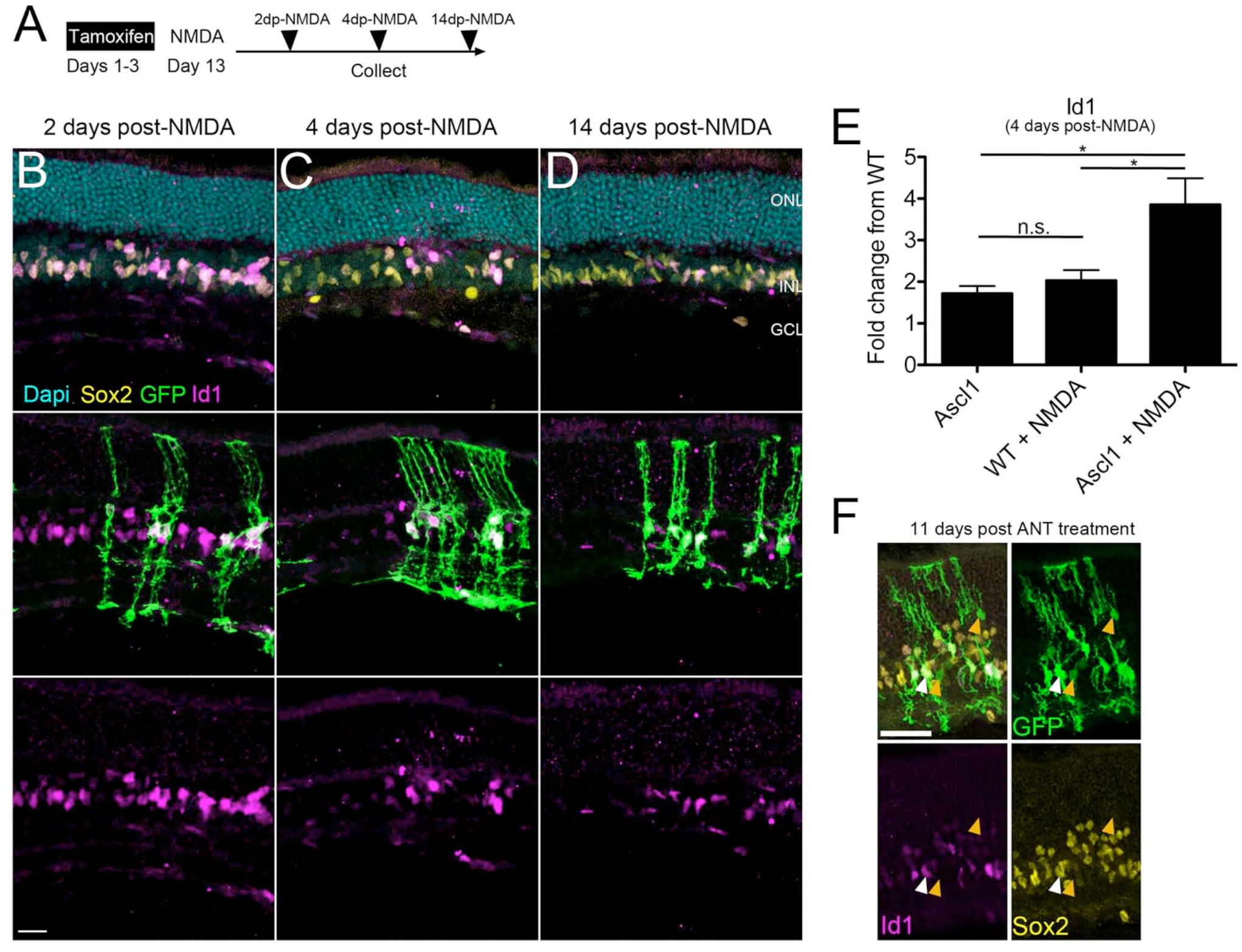
Ascl1-overexpression results in dysregulated STAT-target genes. **A)** Experimental paradigm for analyzing STAT-target genes in Ascl1-overexpressing MG. **B)** Both GFP-Sox2+ and Ascl1-expressing GFP+ MG express Id1 2 days after NMDA damage. **C)** Ascl1+ cells have higher expression of Id1 relative to GFP-MG (Sox2+ cells) 4 days after NMDA damage. **D)** Id1 expression is reduced in GFP-MG but remains in the Ascl1-expressing GFP+ MG 14 days after NMDA damage. Note, the flat Id1+ nuclei in all images not labelled with GFP are endothelial cells. Scale bars for **B-D** are 20 µm. **E)** Graph showing RT-qPCR for Id1 gene expression relative to WT on retinas treated 4 days post-NMDA. One-way ANOVA with Tukey’s post-test; *P < 0.05, N = 4 biological replicates per condition run in triplicate. **F)** Id1 expression is reduced in GFP-MG but remains in the Ascl1-expressing GFP+ MG 14 days after ANT treatment (white arrows). MG-derived neurons (orange arrows) do not express Id1. Scale bar is 40 µm.

## DISCUSSION

The findings in this report highlight several important features of the reprogramming of glia to neuronal progenitors and neurons. The Ascl1 ChIP-seq and DNase-seq data reveal that the reprogrammed MG show similar Ascl1 binding and chromatin accessibility as the newborn mouse RPCs. However, the reprogrammed MG also have new Ascl1 peaks that are not normally found during development in RPCs, sites we label as “inappropriate.” A significant number of these inappropriate Ascl1 binding sites are found at genes that are not relevant to retinal development and are associated with an increase in expression of these genes in the reprogrammed MG. We further identified a consensus DNA binding motif for STAT3 that was significantly associated with these inappropriate Ascl1 peaks. Potential STAT targets that could underlie the effects we observe, include Id1 and Id3, dominant negative regulators of bHLH gene function. Lastly, this study provides the first evidence combining lineage tracing with EdU-labeling to demonstrate new neurons can arise from proliferating MG in an adult mammal.

One potential target of STAT signaling that could limit the regenerative response in mammalian MG is Id1. Following injury to the retina, the STAT target gene Id1 is transiently expressed in MG, but Id1 returns to basal levels within a week. We found that in Ascl1-overexpressing mice, Id1 expression is maintained in the MG, possibly maintaining the cells in a progenitor-like state and preventing them from generating new neurons. ID proteins are known to form dimers with class I bHLH proteins and inhibit their dimerization with class II bHLH factors to inhibit their transcriptional activation activity^35, 36^. Additionally, ID proteins can bind directly with HES proteins to maintain neural progenitors in an undifferentiated state^39–41^. ID proteins have also been shown to play an important role in other organ systems, such as the pancreas, where they can bind Hes1 and participate in the dedifferentiation and fate switching of exocrine cells to endocrine fates^42^. JAK/STAT signaling is known to bias progenitors towards a glial fate in other regions of the nervous system as well^43–45^. For example, inhibition of the JAK/STAT co-receptor gp130 in developing cortical progenitors has been found to increase their neurogenic production at the expense of gliogenesis^46^. Thus, it is possible that STAT activation in astrocytes in other areas of the nervous system may limit their neurogenic potential via ID proteins in a manner similar to that we have described in the retinal MG.

Previous studies have demonstrated the importance of the source cell’s epigenetic landscape and presence of endogenous transcription factors of that source cell during reprogramming^32, 33^. Fibroblast reprogramming to induced neuronal cells show that the newly generated cell types still retain their source cell signature. We previously found this to be the case in MG reprograming as well^16^, where the MG-derived neurons still express low levels of glial genes such as Glul or Aqp4. The present study demonstrates the importance of injury induced STAT pathway activation during reprogramming and shows a successful combinatorial analysis using epigenetic and gene expression datasets to confirm inappropriate transcription factor binding associated with non-productive gene expression.

## ACKNOWLEDGMENTS

The authors acknowledge the following funding sources for supporting this work. Grant #TA-RM-0614-0650-UWA from the Foundation Fighting Blindness to T.A.R., NIH NEI 1R01EY021482 to T.A.R., Allen Distinguished Investigator Award (Paul G. Allen Family Foundation) to T.A.R. and F.R., a NSF Fellowship to M.S.W. (DGE-0718124), the Vision Core Grant P30EY01730, NIH NEI Training Grant (EY07031) to L.T., NIH F32 NRSA to B.A.W. (5 F32 DC016480-02), and a NIH NEI F31 NRSA to N.L.J. (5 F31 EY028412-02). We thank members of the Reh and Bermingham-McDonogh laboratories for their review and valuable discussion regarding the manuscript. We thank the laboratory of C. Trapnell, specifically D. Jackson for her help generating the single-cell RNA-seq data. Lastly, the authors thank M. Nakafuku (Cincinnati Children’s) for the tetO-Ascl1-ires-GFP mice.

## AUTHOR CONTRIBUTIONS

N.L.J. conceived of and performed all in vivo experiments and STAT inhibitor experiments and analyses. L.T. performed EdU-labeling experiments and contributed to the written manuscript. P.N. performed cell culture and WB experiments and contributed to the written manuscript. M.J.H. performed cell culture and contributed to Ascl1 ChIP-seq experiments. B.A.W. assisted in ChIP-seq analyses and the writing of the manuscript. F.R. Performed whole cell electrophysiological recordings. N.R. performed in vivo experiments and contributed to cell counts. M.S.W. conceived of and performed all Ascl1 ChIP-seq and DNase-seq experiments and analyses. A.C. assisted in scRNA-seq analyses. T.A.R. conceived of all experiments and performed analyses on sequencing datasets

## DECLARATION OF INTERESTS

The authors declare no competing interests.

## METHODS

### CONTACT FOR REAGENT AND RESOURCE SHARING

Further information and requests for resources and reagents should be directed to and will be fulfilled by the Lead Contact, Thomas A. Reh.

### EXPERIMENTAL MODEL AND SUBJECT DETAILS

#### Mice

*Glast-CreER:LNL-tTA:tetO-mAscl1-ires-GFP* mice and *rtTA germline*: *tetO-mAscl1-ires-GFP* mice used in this study were from a mixed background of C57BL/6, B6SJLF1, and other backgrounds present at The Jackson Laboratory. Mice of both sexes were used in this study. For in vivo experiments, adult mice over the age of postnatal day 40 were used. Mice were housed in the specific-pathogen-free (SPF) animal facility at the University of Washington, Seattle, WA. Mice were housed under controlled conditions. Mice underwent no previous treatments prior to testing. All procedures performed in this study were approved by the Institutional Animal Care and Use Committee at the University of Washington, Seattle.

#### Primary Cell Culture

Postnatal day 0, 11/12 retinas of both sexes from *rtTA germline:tetO-Ascl1-ires-GFP* mice were digested with papain/DNase to single cells and MG were grown in culture as previously described^34, 47^. In brief, retinas were placed in a papain solution with 180 units/mL DNase (Worthington) and incubated at 37 °C for 10 min. Cells were triturated and added to an equal ovomucoid (Worthington) volume then spun down at 300 g at 4 °C to pellet. Cells were then resuspended in Neurobasal with 10 % FBS (Clontech), mEGF (100 ng/mL; R&D Systems), 1 mM L-glutamine (Invitrogen), N2 (Invitrogen), and 1 % Penicillin-Streptomycin (Invitrogen) with two retinas plated per 10 cm^2^ at 37 °C. Media was changed every 2 days until confluent monolayers of MG were passaged after 7 DIV and doxycycline was added to induce expression of Ascl1.

### METHOD DETAILS

#### Animals

All mice were housed at the University of Washington. All experiments and protocols were approved by the University of Washington’s Institutional Animal Care and Use Committee. Adult mice used for in vivo experiments were *Glast-CreER:LNL-tTA:tetO-mAscl1-ires-GFP* mice and have been previously described^16^. The *Glast-CreER* and *LNL-tTA* mice were from Jackson Labs and the *tetO-mAscl1-ires-GFP* mice were a gift from M. Nakafuku (University of Cincinnati). The *rtTA germline*: *tetO-mAscl1-ires-GFP* mice for in vitro experiments were generated by crossing Nakafuku’s *tetO-mAscl1-ires-GFP* mice onto the germline rtTA mice from Jackson Labs. Mice of both sexes were used in this study and adult mice were treated at ages comparable to our previously described study^16^.

#### Ascl1 Chromatin Immunoprecipitation-Sequencing (ChIP-Seq)

P0 retinas or cultured, post-natal day 12, Müller glia (+/-Ascl1 overexpression, *rtTA germline:tetO-Ascl1-ires-GFP* mice ± doxycycline) were digested with papain/DNase to single cells and fixed with 0.75 % formaldehyde for 10 minutes at room temperature. Sonication was performed with a probe sonicator (Fisher Scientific): 12 pulses, 100 J/pulse, Amplitude: 45, 45 seconds cooling at 4 °C between pulses. Immunoprecipitation performed with 40 μL anti-mouse IgG magnetic beads (Invitrogen Cat: 110.31) and 4 μg mouse anti-MASH1 antibody (BD Pharmingen Cat: 556604) or 4 μg mouse IgG against chromatin from 5 million cells per condition according to Diagenode LowCell Number Kit using IP and Wash buffers as described in^28^. Libraries were prepared with standard Illumina adaptors and sequenced to an approximate depth of 36 million reads each. Sequence reads (36 bp) were mapped to the mouse mm9 genome using bwa (v 0.7.12-r1039). Merging and sorting of sequencing reads from different lanes was performed with SAMtools (v1.2). The HOMER software suite was used to determine and score peak calls (‘findPeaks’ function, v4.7) as well as motif enrichment (‘findMotifs’ function, v4.7, using repeat mask). For STATi and control Ascl1 ChIP-seq, reads were aligned to the mm10 genome using Bowtie2. The .sam files were converted to sorted .bam files using SAMtools. MACS2 was used to call peaks with default settings using the broad peaks annotation. Peak overlap analyses were performed using Bedops. The control Ascl1 ChIP-seq .bam file was downsampled by a factor of 0.69 to normalize the number of mapped reads over the common peaks found between treatment and control samples. This downsampled .bam file was used for all analyses. Differential accessibility analysis in Ascl1 ChIP-seq peaks was determined using edgeR as detailed in the edgeR user guide.

#### DNase I Hypersensititivy-Sequencing (DNase-Seq)

Detailed protocols can be found at encodeproject.org. In brief: nuclei from retina were isolated using 25 strokes of a dounce homogenizer, tight pestle, in 3 mL homogenization buffer (20 mM tricine, 25 mM D-sucrose, 15 mM NaCl, 60 mM KCl, 2 mM MgCl_2_, 0.5 mM spermidine, pH 7.8) and filtered through a 100 μm filter and washed with Buffer A (15 mM Tris-HCl, 15 mM NaCl, 60 mM KCl, 1 mM EDTA, 0.5 mM EGTA, 0.5 mM spermidine). Nuclei from Müller glia were isolated using TrypLE (Thermofisher) to obtain single cells, followed by incubation with 0.04 % IGEPAL in Buffer A for 10 minutes at 4 °C. Nuclei were incubated at 37 °C for 3 minutes in limiting concentrations of DNaseI enzyme in Buffer A with calcium supplement. The reaction was stopped using equal volume of Stop Buffer (50 mM Tris-HCl, 100 mM NaCl, 0.1 % SDS, 100 mM EDTA, 1 mM spermidine, 0.5 spermine pH 8.0) and subsequently treated with proteinase K and RNase A at 55 °C. Small (<750 bp) DNA fragments were isolated by sucrose ultracentrifugation and end repaired and ligated with Illumina compatible adaptors. Sequence reads were mapped to mm9 using bowtie (v 0.12.7) and DNaseI peak calling performed with Hotspot (http://www.uwencode.org/proj/hotspot/).

#### Electrophysiology

Recordings were performed under identical conditions to our previous study^16^. All mice underwent dark-adaptation prior to euthanasia. Retinas were then removed, dissected, embedded in agar, and cut into 200 μm thick slices under infrared visualization. All prep was performed in Ames medium at 32-34 °C and oxygenated with 95 % O_2_/ 5 % CO_2_. Slices were placed under the microscope and perfused with oxygenated Ames at a rate of ∼8 mL per minute. Cells were targeted for recording using video DIC with infrared light (>950 nm), two-photon (λ = 980 nm), or confocal (λ = 488 nm) microscopy. Cells targeted for light responses under infrared conditions were exposed to full-field illumination via blue and green LEDs from a customized condenser. Cells targeted with the 488 nm laser were predominantly used to record *R*_in_ and *V*_rest_ properties. Recordings were performed using pulled glass pipettes (5-6 MΩ) and filled with solution containing (in mM): 123 K-aspartate, 10 HEPES, 1 MgCl_2_, 10 KCl, 1 Cacl_2_, 2 EGTA, 0.5 Tris-GTP, 4 Mg-ATP, and 0.1 Alexa-594 hydrazide.

#### Fluorescence-activated cell sorting (FACS)

After euthanasia of mice, eyes were removed and retina dissected out and isolated from vitreous and retinal pigmented epithelium. Retinas were then dissociated in a Papain and DNase I solution for 20 minutes at 37 °C on a nutator. After incubation, retinas were gently triturated to generate a single-cell suspension, followed by the addition of Ovomucoid. Cells were then spun down at 300 g at 4 °C then resuspended in Neurobasal medium, passed through a 35 μm filter, and transferred to a BSA coated tube prior to FACS-purification. FACS was performed on a BD FACSAria III Cell Sorter (BD Biosciences) and the GFP-positive fraction was collected for single-cell RNA-seq. 44,000 cells in total were captured for single-cell RNA-seq.

#### Immunohistochemistry (IHC)

Upon euthanasia of adult mice, eyes were removed and corneas dissected away. Globes were then fixed for 1 hour in 4 % PFA in PBS followed by an overnight incubation in 30 % sucrose in PBS at 4 °C. Fixed eyes were then frozen in O.T.C. (Sakura Finetek) at −80 °C until sectioned. Frozen eyes were sectioned on a cryostat (Leica) at 16-18 μm thick and were stored at −20 °C until staining. All washes were 3 times 20 minutes in PBS on a rotating plate at room temperature. Slides were washed then blocked in 10 % normal horse serum, 0.5 % Triton X-100 in PBS for a minimum of 1 hour prior to primary. All primary antibodies were diluted in blocking solution and applied to tissue for a minimum of 3 hours. Slides were then washed and incubated with secondary antibodies for a minimum of 2 hours, which were diluted in PBS. Slides were then washed and cover slipped with Fluoromount-G (SouthernBiotech). Primary antibodies used were: rabbit anti-Cabp5 (a gift from F. Haeseleer, 1:500), rabbit anti-PSD95 (Abcam, 1:100, Ab-18258-100), mouse anti-Ctbp2 (BD Biosciences, 1:1000, 612044), chicken anti-GFP (Abcam, 1:500, Ab13970), goat anti-Otx2 (R&D Systems, 1:100, BAF1979), goat anti-Sox2 (Santa Cruz, 1:100, SC-17320), rabbit anti-Opn1sw (Millipore, 1:300, AB5407), rabbit anti-Id1 (Biocheck, 1:1000, BCH-1/#37-2). Secondary antibodies used were all donkey anti-species (Life Technologies) and were diluted 1:300 with a 1:100,000 DAPI (Sigma) in PBS and were applied in dark conditions. TUNEL staining was performed using the Promega TUNEL kit. EdU-labelling was performed using the Thermo Fischer Scientific Click-iT EdU system.

#### Injections

Intraperitoneal injections of tamoxifen were administered to adult mice for up to five consecutive days to induce expression of the *tetO-mAscl1-ires-GFP* gene. Tamoxifen was administered at a concentration of 1.5 mg per 100 μL of corn oil. Intravitreal injections of NMDA were administered at a concentration of 100 mM in PBS at a volume of 1.5 μL. Intravitreal injections of TSA (Sigma) were administered at a concentration of 1 μg per μL in DMSO at a volume of 1.5 μL. Intravitreal injections of SH-4-54 STAT-inhibitor (Selleck Chem) were administered at a concentration of 10 mM in TSA containing DMSO at a volume of 1.5 μL. All intravitreal injections were performed on isoflurane-anesthetized mice using a 32-gauge Hamilton syringe. For EdU-labelling experiments, NMDA and EdU were intravitreally co-injected in a mixture containing 1 μL of 34mM NMDA, 0.5 μL EdU (5 mg/mL), and 0.5 μL PBS at a volume of 2 μL. Two days later the TSA, STATi, and EdU were intravitreally co-injected in a mixture containing 1 μL TSA (2 mg/mL), 0.5 μL EdU (5 mg/mL), and 0.5 μL SH-4-54 (25 mg/mL) at a volume of 2 μL. From treatment days 9 through 12, EdU was intraperitoneally injected twice daily for a total of 8 injections at a concentration of 1 mg/mL EdU at a volume of 100uL in PBS.

#### Microscopy/cell counts

All images were taken on a Zeiss LSM880 confocal microscope. For cell counts, all images were taken at the same magnification and a minimum of 4 fields per retina were captured as a Z-stack then analyzed in ImageJ (NIH, Bethesda, MD). To be counted as a positive cell, the marker of interest needed to be viewed in at least 3 planes of the Z-stack to ensure accurate co-localization. Counts were summed together for a single retina then percent co-localization was calculated per retina. For the Id1, Sox2, GFP staining figure, all images were captured at the same magnification, laser settings, and detector settings. In addition, all sections were stained at the same time in the same manner to show comparable relative protein expression in each treatment. For synaptic staining images, the Airyscan detector on the Zeiss LSM880 was used to capture Z-stacks and maximize resolution and co-localization of synaptic proteins. Airyscan images were then processed in Amira image software (FEI). The GFP channel was masked and Psd95 that was within this GFP mask in the OPL was displayed to highlight Müller glial-derived neurons synaptic input. Similarly, the GFP channel was masked and Ctbp2 that was within this GFP mask in the IPL was displayed to highlight Müller glial-derived neurons synaptic output.

#### Quantitative reverse transcription PCR (RT-qPCR)

*Glast-CreER:LNL-tTA:tetO-mAscl1-ires-GFP* mice received 3 days of tamoxifen injections to induce Ascl1 expression on treatment days 1 through 3. On treatment day 13, WT and Ascl1 expressing mice received an intravitreal injection of 100 mM NMDA in 1.5 μL volume. Four days after NMDA administration WT and Ascl1 expressing mice were euthanized and retinas collected and digested in TRIzol reagent (Thermo Fischer). RNA was extracted and collected in miRNeasy Mini Kit columns in accordance with manufacturer instructions (Qiagen). Reverse transcription was performed on 1 μg of purified RNA using the iScript Reverse Transcription Supermix kit (Bio-Rad). The cDNA was then added to SsoFast (Bio-Rad) for qPCR. A total of 4 biological replicates were run for each condition (WT, WT + NMDA, Ascl1, Ascl1 + NMDA) and each biological replicate was run in triplicate. Id1 cycles were subtracted from housekeeping gene Gapdh (ΔCt) and then subtracted from WT (ΔΔCt) to determine fold change 2^(–_ΔΔ_Ct)^. Primers for Gapdh were (5′-GGCATTGCTCTCAATGACAA-3′ and 5′-CTTGCTCAGTGTCCTTGCTG-3′) and primers used for Id1 were (5′-TACGACATGAACGGCTGCTACTCA-3′ and 5′-TTACATGCTGCAGGATCTCCACCT-3′).

#### Single-cell RNA-sequencing (Single-cell RNA-seq)

Four ANTSi-treated mice (8 eyes) had their retinas pooled for FACS-purified cells and were spun down at 300 g at 4 °C then resuspended at a concentration of 1000 cells per μL in a 0.04 % BSA in PBS solution. Cells were processed through the 10x Genomics Single Cell 3’ Chip and processed through the standard Chromium Single Cell 3’ Reagent Kits User Guide protocol with a target capture of 4,000 cells. Library QC was determined by Bioanalyzer (Agilent). Libraries were then sequenced on Illumina NextSeq 500/550 vs kit and reads were processed through 10x Genomics Cell Ranger pipeline. Reads were aligned to the mm10 genome and the filtered output files from Cell Ranger were processed in R using tools in the Seurat package. For Seurat analyses, the newly generated single-cell RNA-seq dataset from ANTSi-treated mice were compared with our previously generated WT and ANT-treated datasets^16^.

#### Western Blot/Analysis

Retinas from P11 *rtTA germline*: *tetO-mAscl1-ires-GFP* mice were dissociated and MG were grown in culture as previously described^34, 47^. Confluent monolayers of MG were passaged and doxycycline (1:500) was added to the media to overexpress Ascl1 for 6 days, and a subset of cultures received SH-4-54 STAT-inhibitor (Selleck Chem) on the fifth day of treatment for 24 hours. MG cultures were lysed with buffer containing 25 mM Tris-HCl pH 7.5, 150 mM NaCl, 1 mM EDTA, 1 % Triton X-100, 5 % glycerol, 1X protease inhibitor cocktail, and 1X phosphatase inhibitor cocktail and equal amounts of protein samples were loaded and run in a 4 % to 20 % SDS gel (Bio-Rad Laboratories). Protein was transferred to a polyvinylidene fluoride membrane (Thermo Fisher Scientific, Waltham, MA, USA), blocked (5 % BSA and 0.1 % Tween 20 in 1X TBS) for at least 1 hour at room temperature and stained with primary antibodies (Phospho-STAT3 Y705 and STAT3, R&D Systems 9131S and 9132S) diluted in blocking solution overnight at 4 °C. Membranes were washed with 0.1 % Tween 20 in 1X TBS and then incubated with HRP-conjugated secondaries (Bio-Rad Laboratories) diluted in blocking solution for 1 hour at room temperature. Signals were visualized on X-ray film with a commercial substrate (SuperSignal West Dura Extended Duration Substrate; Thermo Fisher Scientific) and quantified using ImageJ software.

### QUANTIFICATION AND STATISTICAL ANALYSIS

#### Ascl1 ChIP-Seq

The HOMER software suite was used to determine and score peak calls (‘findPeaks’ function, v4.7) as well as motif enrichment (‘findMotifs’ function, v4.7, using repeat mask). Peak overlap analyses were performed using Bedops. Gene Ontology analyses of Ascl1 ChIP-seq peaks were generated using GREAT algorithm.

#### Immunohistochemistry

For Otx2 quantification, retinas from 16 ANT-treated mice and 13 ANTSi-treated mice were analyzed. For Cabp5 quantification, retinas from 6 ANT-treated mice and 8 ANTSi-treated mice were analyzed. An unpaired *t*-test was performed using Graphpad Prism for each figure. Data is presented as mean ± SEM.

#### RT-qPCR

Id1 expression was performed on 4 biological replicates from each condition.

#### Single-cell RNA-seq

Quantification of the percent cells in each cluster was performed in R using the Seurat toolkit. Cells that were located in either the Müller glial, Progenitor-like, or MG-derived neuron clusters were identified by which treatment they came from (WT, ANT, ANTSi). The total number of cells in each cluster from each treatment was then divided by the total number of cells from that corresponding treatment to generate percentages, which are shown in the bar graph. The heatmap was created using the doHeatmap function from the Seurat toolkit in R. The intensity of the log2 expression is shown.

#### Western Blot

Quantification of gel bands was performed using ImageJ.

### DATA AND SOFTWARE AVAILABILITY

Database/accession numbers for Ascl1-ChIP-seq and single-cell RNA-seq datasets will be deposited and described here upon acceptance of this manuscript.

---

**Figure S1.**
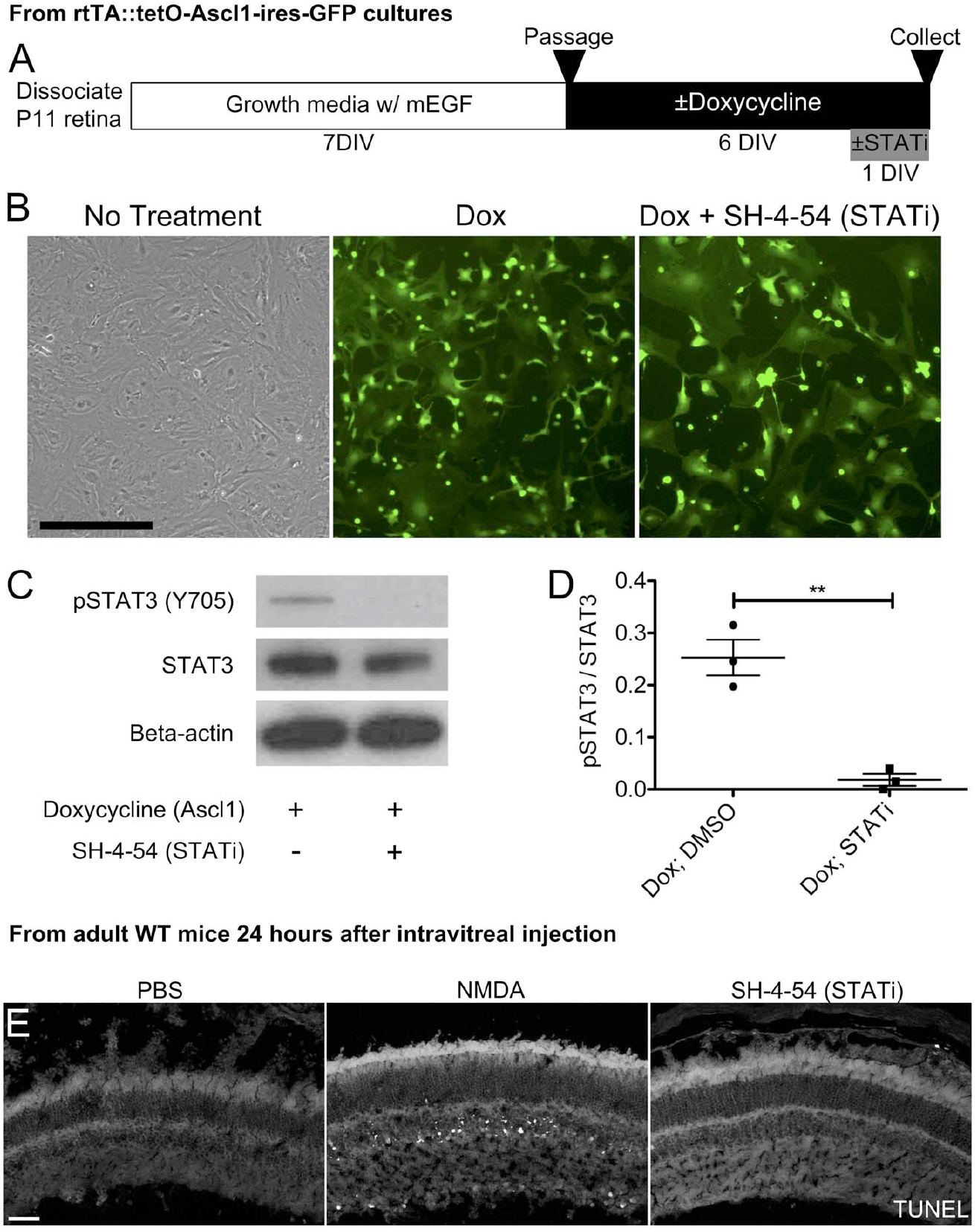
STAT inhibitor SH-4-54 inhibits pSTAT3 in MG. **A)** Experimental paradigm for analyzing STAT3 phosphorylation by western blot in MG. P11 retinas were dissociated and grown in a growth media containing mEGF for 7 DIV. MG were passaged to eliminate residual neurons and cultured in a media containing doxycycline to induce Ascl1 expression for 6 DIV. On the sixth day, the STATi SH-4-54 was added then cells were harvested 24 hours later for WB. **B)** Representative images from confluent cultures of non-treated, Dox-treated, and Dox + STATi-treated MG just before harvesting. **C)** Representative western blot for pSTAT3 (Y705), STAT3, and Beta-actin on non-treated, Dox-treated, and Dox+ STATi-treated MG cultures. **D)** Graph showing quantification of western blot bands was significantly different by One-way ANOVA with Tukey’s post-test, *P < 0.05, ***P < 0.0001. **E)** Representative image of WT retinas injected with either PBS, NMDA, or SH-4-54 and collected 24 hours later for TUNEL staining. Scale bars for **B** and **E** are 25 and 40 µm, respectively.

**Figure S2.**
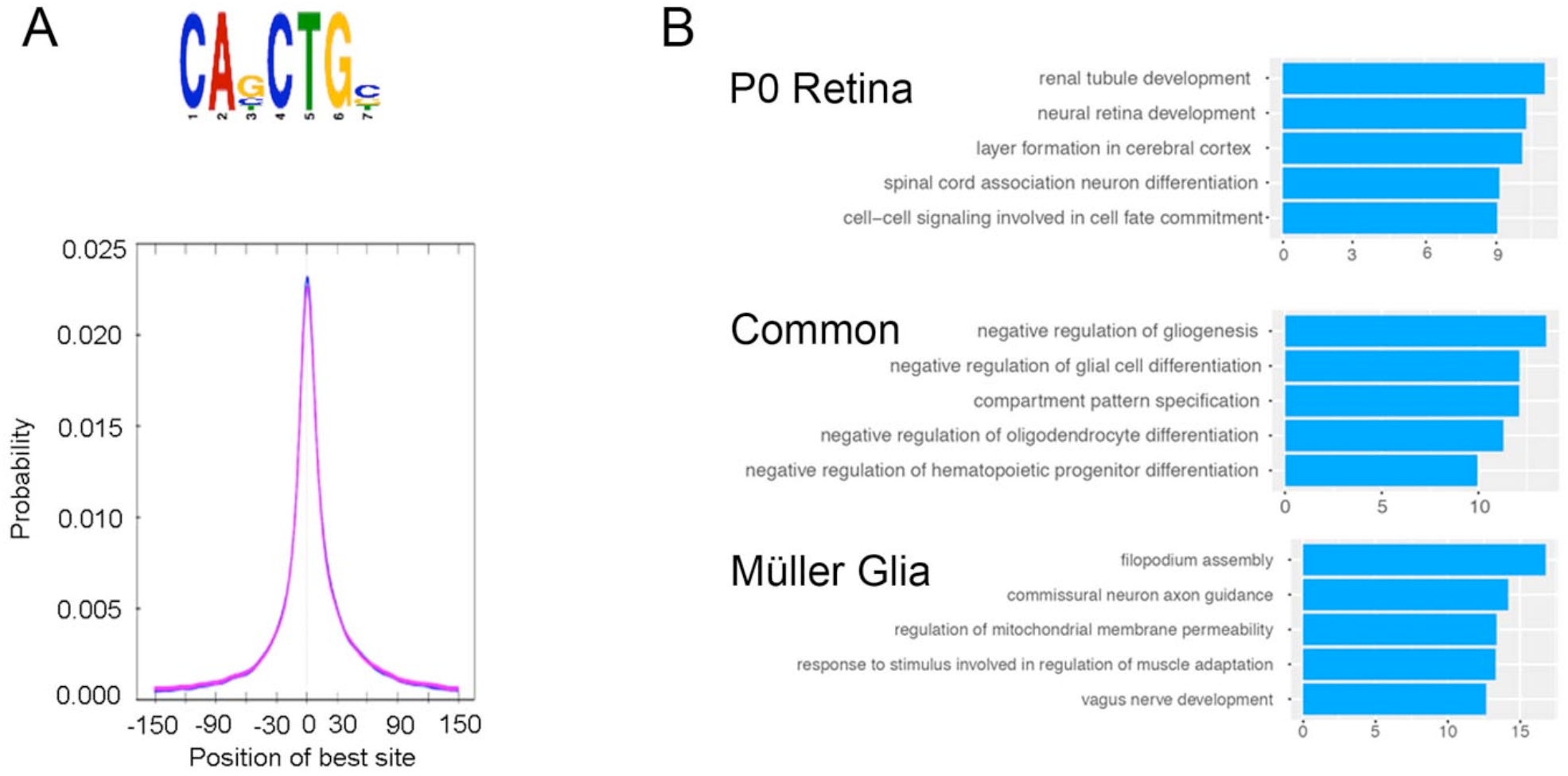
Analysis of P0 and MG Ascl1 ChIP-seq peaks. **A)** Motif enrichment analysis from MEME of top scoring motif from P0 Ascl1 ChIP-seq (2 replicates) and diagram showing central enrichment around the Ebox motif. **B)** Gene ontology analysis from GREAT showing top 5 enriched categories by -log10 (binomial p value) for P0-specific peaks, Common peaks, and MG-specific peaks.

**Figure S3.**
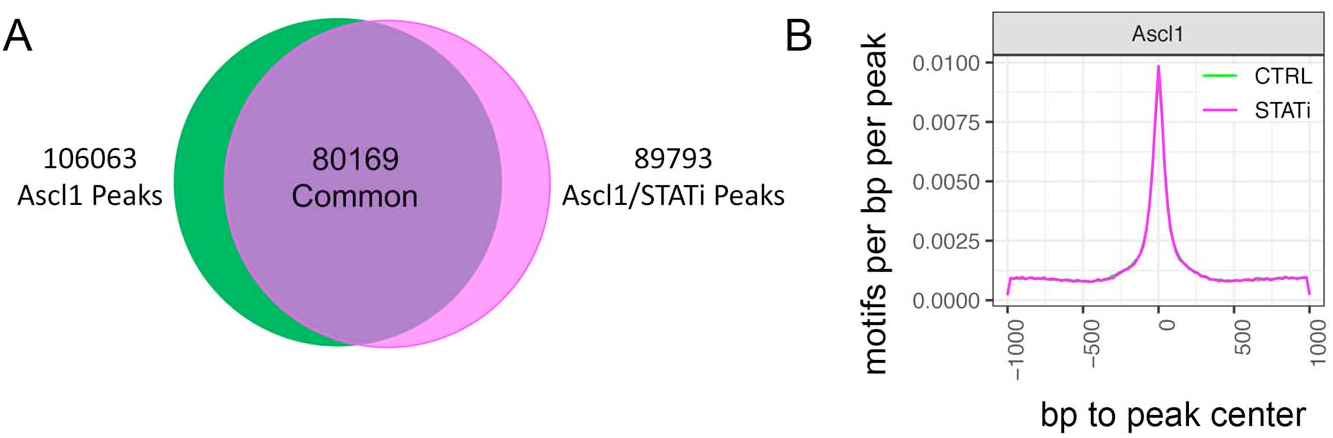
Analysis of MG Ascl1 ChIP-seq peaks from control and STATi treatments. **A)** Venn diagram of all Ascl1 ChIP-seq peaks from control (Ascl1) and STATi (Ascl1/STATi) treatment conditions. 80,169 peaks were common between the two samples. **B)** Diagram from HOMER analysis showing central enrichment around the Ebox motif for control and STATi treatment conditions.

**Figure S4.**
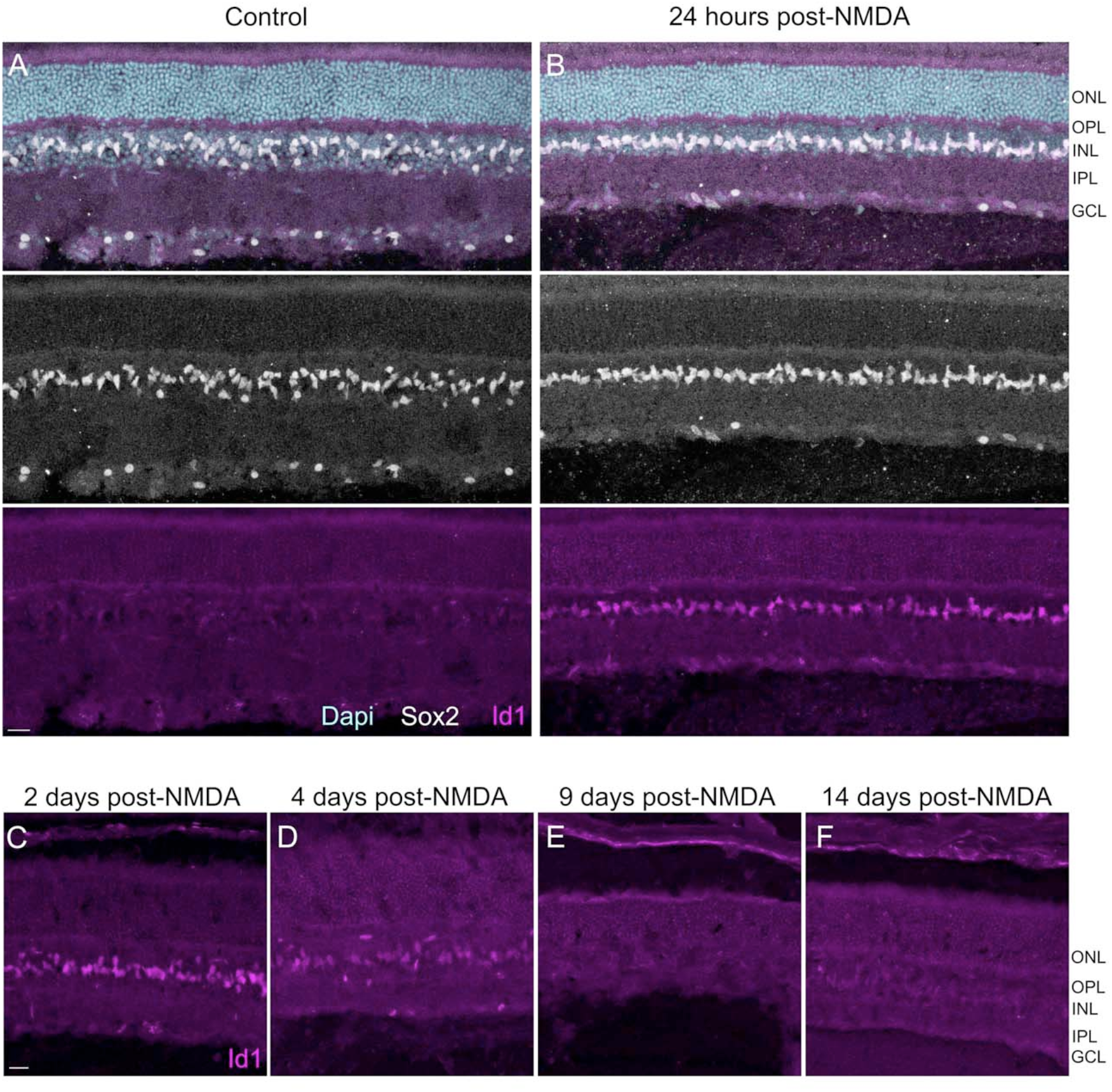
Stat pathway is transiently activated during NMDA damage. **A)** Example image of WT retina stained for MG (Sox2) and Stat pathway activation (Id1). **B)** WT retina 24 hours after NMDA treatment shows Id1 expression in all MG. **C-F)** Id1 expression begins to decrease at 4 days post-NMDA and is undetectable by 9 days. Scale bars are 20 µm.

